# Reversal of an existing hearing loss in *Spns2* mutant mice

**DOI:** 10.1101/2023.05.02.539081

**Authors:** Elisa Martelletti, Neil J. Ingham, Karen P. Steel

## Abstract

Hearing loss is highly heterogeneous but one common form involves a failure to maintain the local ionic environment of the sensory hair cells reflected in a reduced endocochlear potential. We used a genetic approach to ask if this type of pathology can be reversed, using the *Spns2*^*tm1a*^ mouse mutant known to show this defect. By activating *Spns2* gene transcription at different ages after the onset of hearing loss we found that an existing auditory impairment can be reversed to give close to normal thresholds for an Auditory Brainstem Response (ABR), at least at low to mid stimulus frequencies. Delaying the activation of *Spns2* led to less effective recovery of ABR thresholds suggesting there is a critical period for intervention. Early activation of *Spns2* not only led to improvement in auditory function but also to protection of sensory hair cells from secondary degeneration. The genetic approach we have used to establish that this type of hearing loss is in principle reversible could be extended to many other diseases using available mouse resources.

**Significance statement:** Neurological diseases are often thought to be irreversible, including hearing loss. In this study we found that one type of hearing loss can be reversed as long as the treatment is delivered within a critical period early in disease progression. This result is a proof-of-concept that hearing loss not only can be avoided but also may be reversed. This genetic approach can be used for a wide range of diseases using existing mouse resources.

---

Hearing impairment is very common in the population and can begin at any age. Over half of adults in their 70s have a significant hearing loss. Hearing impairment isolates people from society, can be associated with depression and cognitive decline, and is a major predictor of dementia (Fellinger et al 2012; Karpa et al 2010; Mick et al 2014; Livingston et al 2017). The only remedies currently available are devices such as hearing aids and cochlear implants, but these do not restore normal function. There is a large unmet need for medical approaches to slow down or reverse hearing loss.

Treatment strategies for genetic deafness are being developed using the mouse, including gene suppression using siRNAs, antisense oligonucleotides to correct splicing, gene replacement, and gene editing to repair single base mutations (Yeh et al 2020; Gao et al 2018; Lentz et al 2013; Chang et al 2015; Al-Moyed et al 2015; György et al 2019; Isgrig et al 2017; Choi et al 1011; Ding et al 2020; Omichi et al 2019; Taiber & Avraham 2019). These studies usually involve introduction of the agent into the mouse inner ear soon after birth when the auditory system is immature, at a stage corresponding to around 16 to 24 weeks of gestation in humans, making direct translation challenging. One report suggests that introduction of otoferlin (*Otof*) sequences into the mouse inner ear at later stages leads to improved hearing in *Otof* mutants that would otherwise show abnormal inner hair cell synaptic function and deafness (Akil et al 2019). We also are interested in asking if an existing hearing impairment can be reversed, because the main demand for treatments is from people who already have hearing loss, particularly progessive age-related hearing loss.

One significant pathological type underlying age-related hearing loss involves the maintenance of the endolymph, the high-potassium, low-sodium fluid that bathes the upper surface of sensory hair cells. In the cochlea, endolymph is maintained at a positive resting potential, the endocochlear potential (EP), of around +120mV which is essential for normal hair cell sensitivity. We have used a genetic approach in *Spns2* mutant mice to investigate the possibility of reversing hearing loss linked to an EP deficit.

## Results

### Reversal of hearing loss in *Spns2* mutants

*Spns2* encodes Spinster homolog 2, a sphingosine-1-phosphate (S1P) transporter. *Spns2* mutant mice show a rapidly-progressive hearing loss associated with a dramatic decline in EP between two and three weeks after birth (Chen et al 2014; Fig 1D). As EP develops to high levels at first in *Spns2* mutants (Fig 1D; Chen et al 2014), we considered ways of restoring it to normal levels after the onset of hearing loss. The *Spns2* mutation used is the *Spns2*^*tm1a*^ mutation, a knockout-first, conditional-ready design that has a large DNA construct inserted into an intron that disrupts gene expression (Fig 1A) (Skarnes et al 2011). The construct is flanked by FRT sites that recombine on exposure to Flp recombinase, removing the construct and restoring gene function (Chen et al 2014).

**Figure 1.**
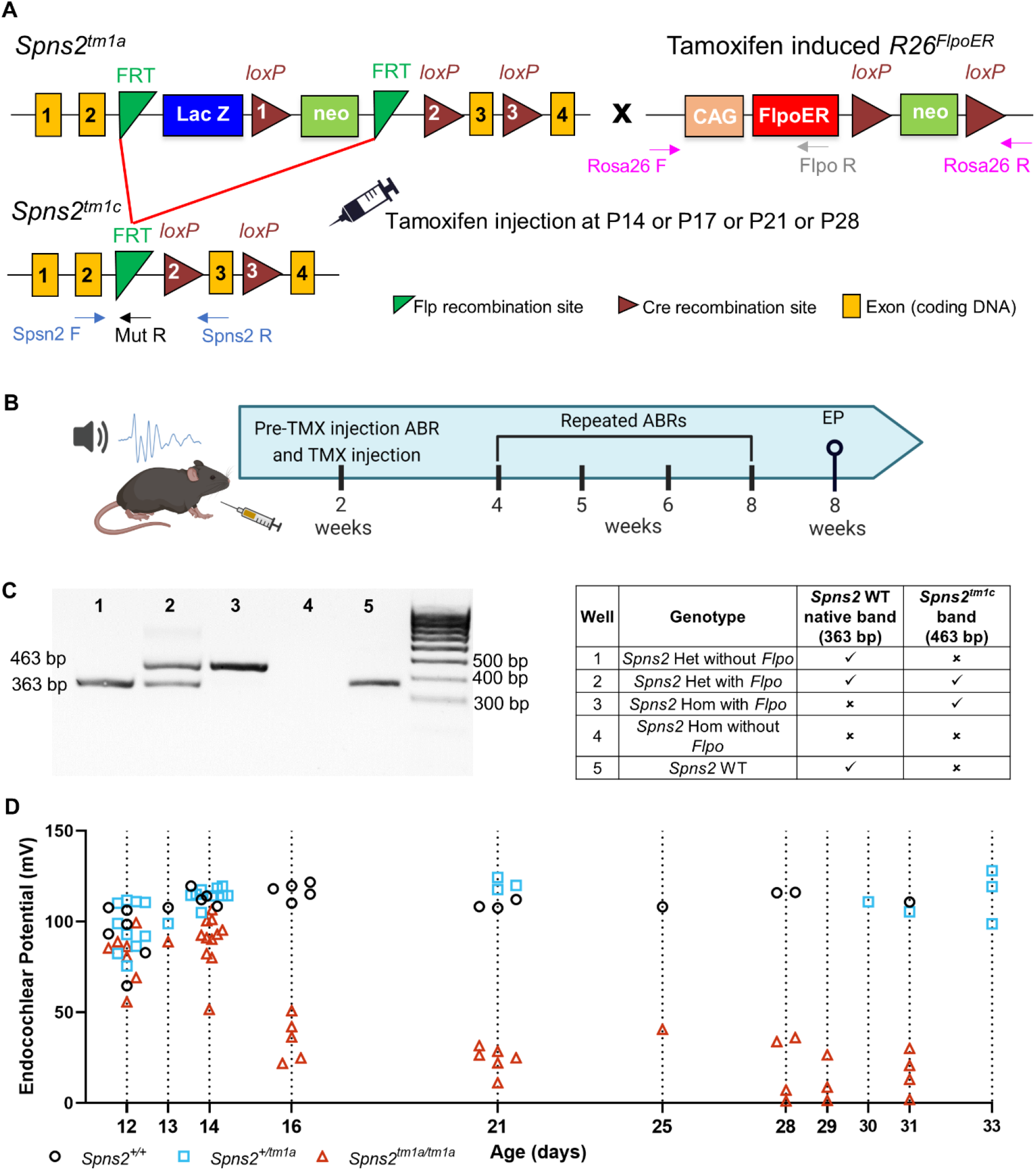
**A**. Diagram showing the design of the *Spns2*^*tm1a*^ and *Spns2*^*tm1c*^ alleles. Yellow boxes show exons, green triangles show FRT sites, brown triangles show loxP sites, blue and green boxes show the neomycin resistance and *LacZ* genes, and arrows marked Fw and Rv indicate the locations of the genotyping primer sites. Upon tamoxifen injection, in the mice carrying the *R26*^*FlpoER*^ allele, the Flpo recombinase induces recombination between the FRT sites and the mutagenic cassette is deleted, resulting in the generation of the *Spns2*^*tm1c*^ allele which is functional (Chen et al 2014). **B**. Diagram showing the experimental design of this study, using 2 weeks as the starting time-point as an example. Auditory Brainstem Responses (ABRs) were recorded at P14, P17, P21, or P28 prior to intraperitoneal tamoxifen injection (TMX, dose 0.2mg/g). Repeated ABR tests were performed up to 8 weeks old when the endocochlear potential (EP) was also measured as a terminal procedure. Diagram generated in Biorender. **C**. Agarose gel showing the PCR product of the *Spns2* Fw and Rv primers (panel A) from pinna samples after tamoxifen injection. The native wildtype (WT) *Spns2* allele has a 363bp PCR product, whereas the *Spns2*^*tm1c*^ allele had a 463bp PCR product, which is 100bp larger due to the presence of one FRT and one loxP sites. *Spns2* homozygotes (mutants) with *Flpo* are shown in wells 1 and 5; the recombination of FRT sites caused by Flpo activity results in a single 463 bp band. *Spns2* heterozygotes without and with *Flpo* are found in well 2 and 4, respectively; in well 2, there is only the 363bp *Spns2 WT* native band, whereas in well 4 there are both the 363 bp *Spns2* WT native band and the 463 bp *Spns2*^*tm1c*^ band. In well 3, *Spns2* homozygotes without *Flpo* show no band; there is no PCR product using those Fw and Rv primers because the presence of the large DNA cassette separates the primer binding sites too far to support normal short-range PCR. The *Spns2* wildtype is shown in well 6. **D**. The EP was measured at different ages between P12 and P33 in controls (black circles), *Spns2*^*+/tm1a*^ (cyan squares) and *Spns2*^*tm1a/tm1a*^ (red triangles) mice; individual mouse measurements are plotted as clusters around the dotted line indicating the age. The EP develops to high levels at first in *Spns2*^*tm1a/tm1a*^ mice, but between P14 and P16 the EP drops and then remains at a relatively stable low level. The n of control, *Spns2*^*+/tm1a*^ and *Spns2*^*tm1a/tm1a*^ mice are respectively: at P12 six, ten and seven; at P13 one each; at P14 four, nine and eleven; at P16 five for controls and *Spns2*^*tm1a/tm1a*^; at P21 three, three and six; at P25 one for controls and *Spns2*^*tm1a/tm1a*^; at P28-P33 four, five and eleven. The datasets at P14, P21 and between P28 and P33 days were previously published by our group (Chen et al. 2014).

In the current study, we used a tamoxifen-inducible Flp recombinase encoded by *R26*^*FlpoER*^ (*Flpo*; Lao et al 2012) to restore *Spns2* function at different stages in the disease progression. Recombination was verified in pinna skin by PCR using the same primers that were used for genotyping (Fig 1A, B). A single intraperitoneal tamoxifen injection was given either at postnatal age (P) 14, 17, 21 or 28, spanning the ages immediately before (P14) and after the drop in EP (P16 onwards; Fig 1D). Entire litters were injected, including *Spns2*^*tm1a*^ homozygotes, heterozygotes and wildtypes both with and without *Flpo*.

Prior to tamoxifen injection, auditory function was assessed using Auditory Brainstem Response (ABR) measurements and these recordings were repeated in each mouse at intervals after tamoxifen exposure until the mouse was 8 weeks old, when terminal ABR and EP measurements were collected (Fig. 1B). Control littermates (*Spns2*^*tm1a*^ heterozygotes or wildtypes with or without *Flpo*; Fig.2, black) showed continued maturation of their thresholds from 2 to 5 weeks old (Fig.2A-C), suggesting no adverse impact of tamoxifen injection. *Spns2*^*tm1a*^ homozygotes that did not carry *Flpo* showed progressive worsening of their ABR thresholds with age (Fig.2; red). In contrast, *Spns2*^*tm1a*^ homozygotes carrying *Flpo* showed gradual improvement in their thresholds with age (Fig. 2; teal), indicating reversal of their hearing loss.

**Figure 2.**
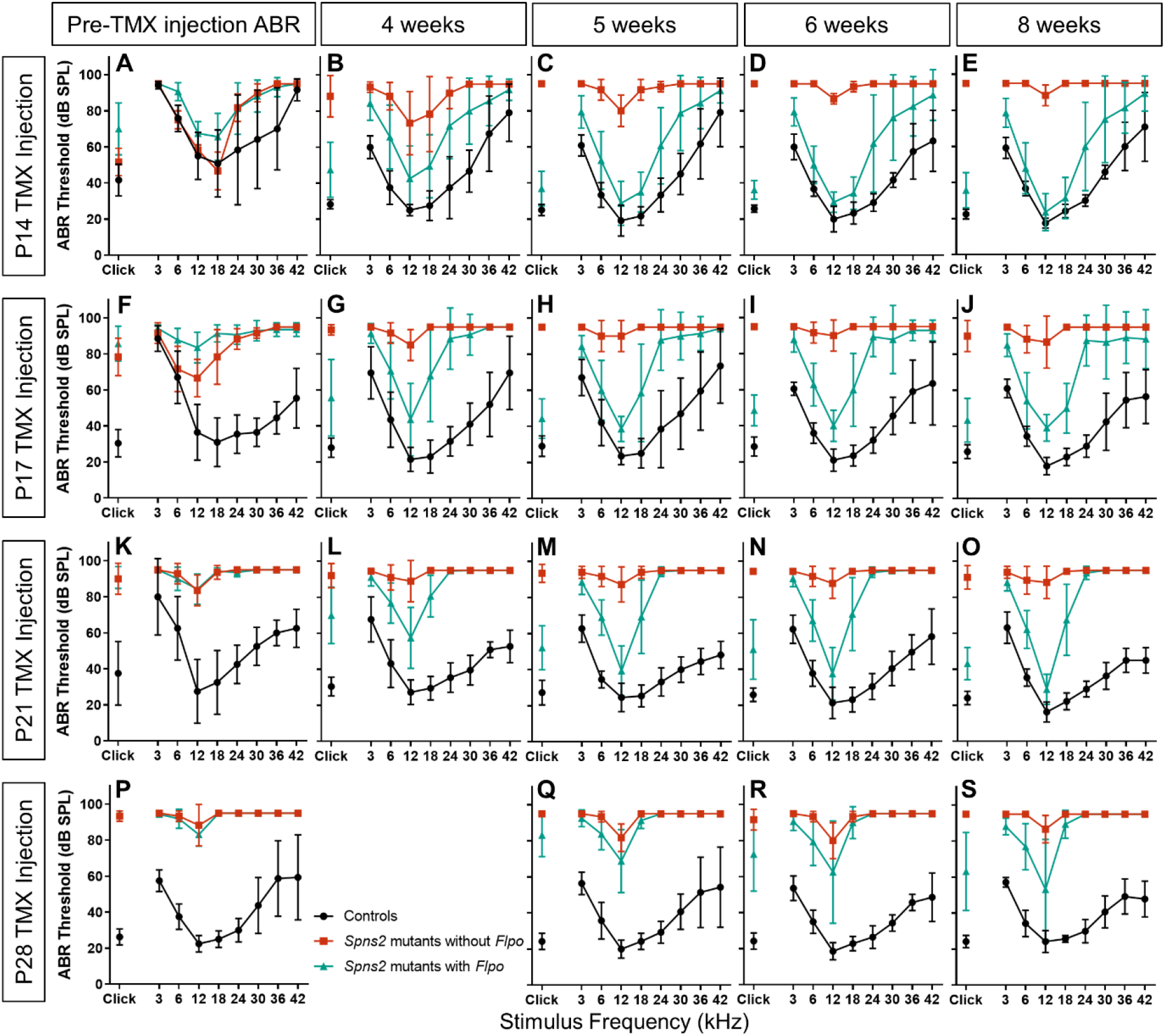
**A-E**. ABR thresholds prior to tamoxifen (TMX) injection at P14 (A), and following at 4, 5, 6 and 8 weeks (respectively B, C, D and E). *Spns2* mutants without *Flpo*, are plotted in red (n = 3), *Spns2* mutants with *Flpo*, are plotted in teal (n = 7), and their control littermates are plotted in black (n = 6 including *Spns2*^*+/+*^ and *Spns2*^*+/tm1a*^ mice with and without *Flpo*). *Spns2* mutants with *Flpo* show a gradual recovery in ABR thresholds at 3, 6, 12, 18 and 24 kHz in comparison with littermate *Spns2* mutants without *Flpo* which have increasing ABR thresholds across all the frequencies. **F-J** ABR thresholds before tamoxifen injection at P17 (F), and after at 4, 5, 6 and 8 weeks (respectively G, H, I and J). *Spns2* mutants with *Flpo* (teal) have higher ABR thresholds at P17 than littermate controls, but after tamoxifen injection, the ABR thresholds at 6, 12 and 18 kHz improve. Controls n=10, *Spns2* mutants without *Flpo* n = 3, *Spns2* mutants with *Flpo* n = 6. **K-O** ABR thresholds before tamoxifen injection at P21 (K), and after at 4, 5, 6 and 8 weeks (respectively L, M, N and O). Only the ABR thresholds at 12 kHz recovered in the *Spns2* mutants with *Flpo* injected with tamoxifen at P21. There is some variability in the recovery of *Spns2* mutants with *Flpo* at 18 kHz. At P21 controls n = 2, *Spns2* mutants without *Flpo* n = 7, *Spns2* mutants with *Flpo* n = 7; whereas at 4, 5, 6 and 8 weeks controls n =11, *Spns2* mutants without *Flpo* n = 9, *Spns2* mutants with *Flpo* n = 14. **P-S** ABR thresholds before tamoxifen injection at P28 (P), and after at 5, 6 and 8 weeks (respectively Q, R and S). The ABR threshold at 12 kHz was 95 dB SPL in two *Spns2* mutants with *Flpo*, but it ranged between 25 and 60 dB SPL in the other six animals. Controls n = 7, *Spns2* mutants without *Flpo* n = 3, *Spns2* mutants with *Flpo* n = 8. Data plotted as mean ± standard deviation.

### A critical period for reversal of hearing loss

By comparing pre and post-tamoxifen ABR thresholds in the same mouse, we observed that injection of tamoxifen at P14 led to the development of near-normal thresholds at frequencies below 24kHz in *Spns2* mutant homozygotes carrying *Flpo* (Fig. 2 A-E). At P17, *Spns2* mutants with *Flpo* already show raised ABR thresholds (Fig. 2 F), but after tamoxifen injection, this hearing loss was reversed at 6, 12 and 18 kHz to near-normal thresholds (Fig. 2F-J). P21 and P28 tamoxifen injections were too late to improve thresholds for frequencies of 18kHz or over, but for 12kHz some improvement was found with injection as late as P28 (Fig. 2K-S). Comparison of the final ABR at 8 weeks old of the 4 injected groups (Fig. 2 E, J, O, S) showed that the earlier the activation of the *Spns2* gene, the more effective was the reversal of the hearing impairment.

Individual *Spns2* mutants with *Flpo* show raised thresholds prior to tamoxifen injection but then almost all of them show a continuing improvement in ABR thresholds with age at the most sensitive frequency, 12 kHz, as indicated by the downward slopes of thresholds plotted in Fig.3A-D. The reversal of hearing impairment was generally stable up to 8 weeks old when the last ABR test was performed. ABR waveforms in *Spns2* mutants with *Flpo* are very similar to those from control littermates in the group injected at P14 suggesting effective restoration of brainstem auditory function (Fig. 3E), although reduced amplitudes and prolonged latencies of individual components of the waveform were observed for later ages of injection (Fig. 3G-I).

**Figure 3.**
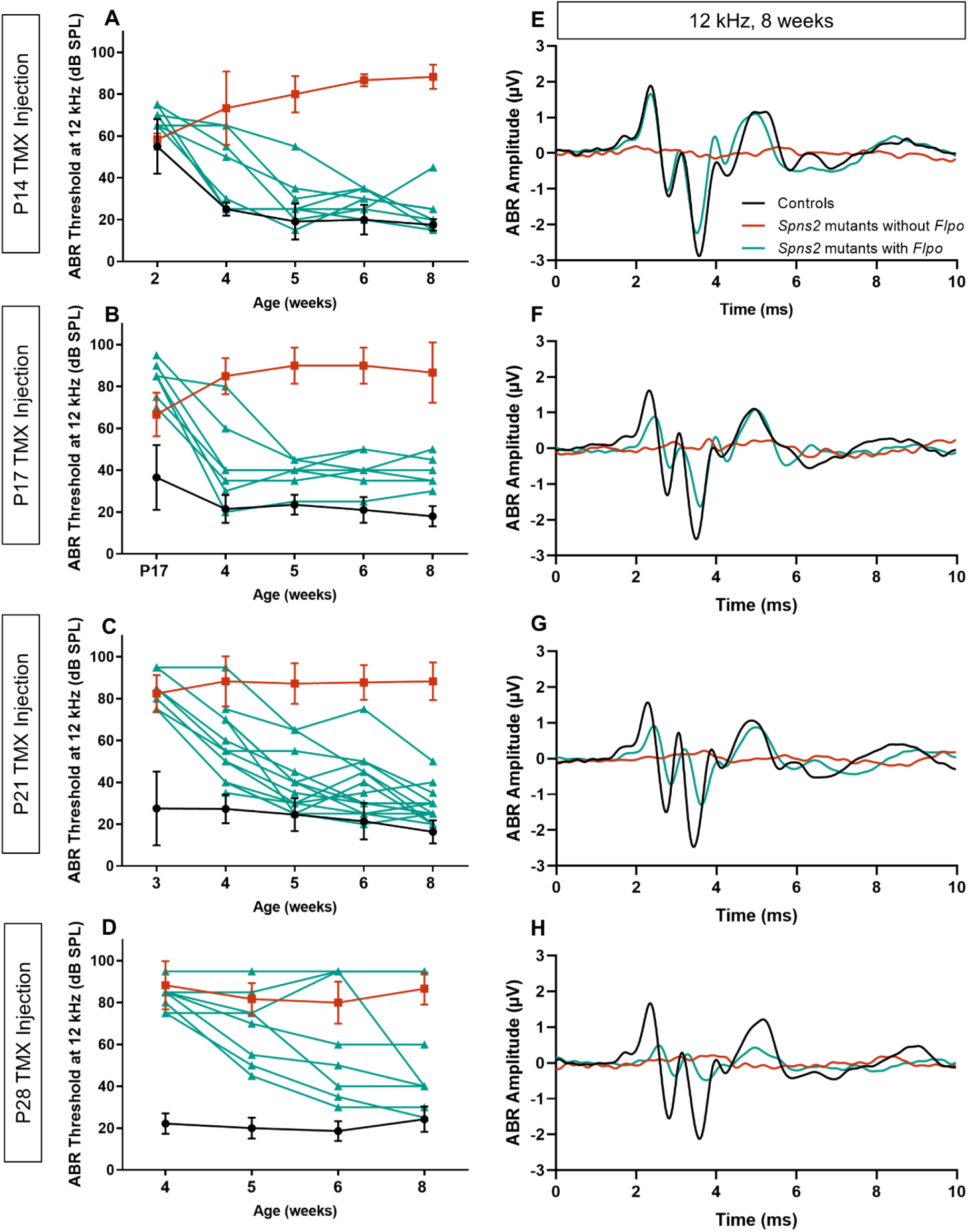
**A-D**. The ABR thresholds at 12 kHz, the best-recovered frequency, are plotted against the ages when the ABRs were tested. At all four injection ages (P14 in A, P17 in B, P21 in C and P28 in D), *Spns2* mutants with *Flpo* have lower ABR thresholds after tamoxifen exposure. There is some variability in *Spns2* mutants with *Flpo* injected at P28, as two out of eight did not recover the ABR threshold at 12 kHz. Controls and *Spns2* mutants without *Flpo* are plotted as mean ± standard deviation in black and red respectively, whereas *Spns2* mutants with *Flpo* are plotted as single animals in teal. For each injected age, the number of the animals is the same as reported in figure 2. **E-I**. The mean evoked waveform at 12kHz, 80dB SPL are plotted for controls (black), *Spns2* mutants without *Flpo* (red) and Spns2 mutants with *Flpo* (teal) aged 8 weeks. Comparable amplitude and latency are observed between controls and *Spns2* mutants with *Flpo* in P14 injected mice. For mice injected at other ages, both reduced amplitude and longer latency are observed in the *Spns2* mutants with *Flpo*, but not as severe as the *Spns2* mutants without *Flpo*. In panel E controls = 6, *Spns2* mutants without *Flpo* = 3 and *Spns2* mutants with *Flpo* = 7; in panel F controls = 10, *Spns2* mutants without *Flpo* = 3 and *Spns2* mutants with *Flpo* = 6; in panel G controls = 11, *Spns2* mutants without *Flpo* = 9 and *Spns2* mutants with *Flpo* = 14; in panel H controls = 7, *Spns2* mutants without *Flpo* = 3 and *Spns2* mutants with *Flpo* = 8.

### Rescue of endocochlear potentials and stria vascularis structure

Measuring EP is a terminal procedure so we cannot track EP recovery directly, unlike tracking hearing via ABR recording. Therefore we measured EP at a single timepoint after the final ABR recording at 8 weeks old in the tamoxifen-injected mice. EP in the control group was around +120mV, a normal level for the mouse (Fig. 4A-D; black). In *Spns2*^*tm1a*^ homozygotes without *Flpo* (red), EP was severely reduced, similar to previously-reported levels in these mutants (Chen et al 2014). The EPs of *Spns2* mutants with *Flpo* (teal) injected at P14 or P17 were higher than those of *Spns2* mutants without *Flpo*, in some cases within the normal range (Fig. 4 A, B). Injection at P21 or P28 led to lower EP measurements than earlier injection (Fig. 4C, D). When EP measurements at all ages of injection and all genotypes were compared with the ABR thresholds at 12 kHz of the same mice, all at 8 weeks old, we found a correlation between lower ABR thresholds and higher endocochlear potential with an R^2^ value of 0.7438 (Fig. 4E).

**Figure 4.**
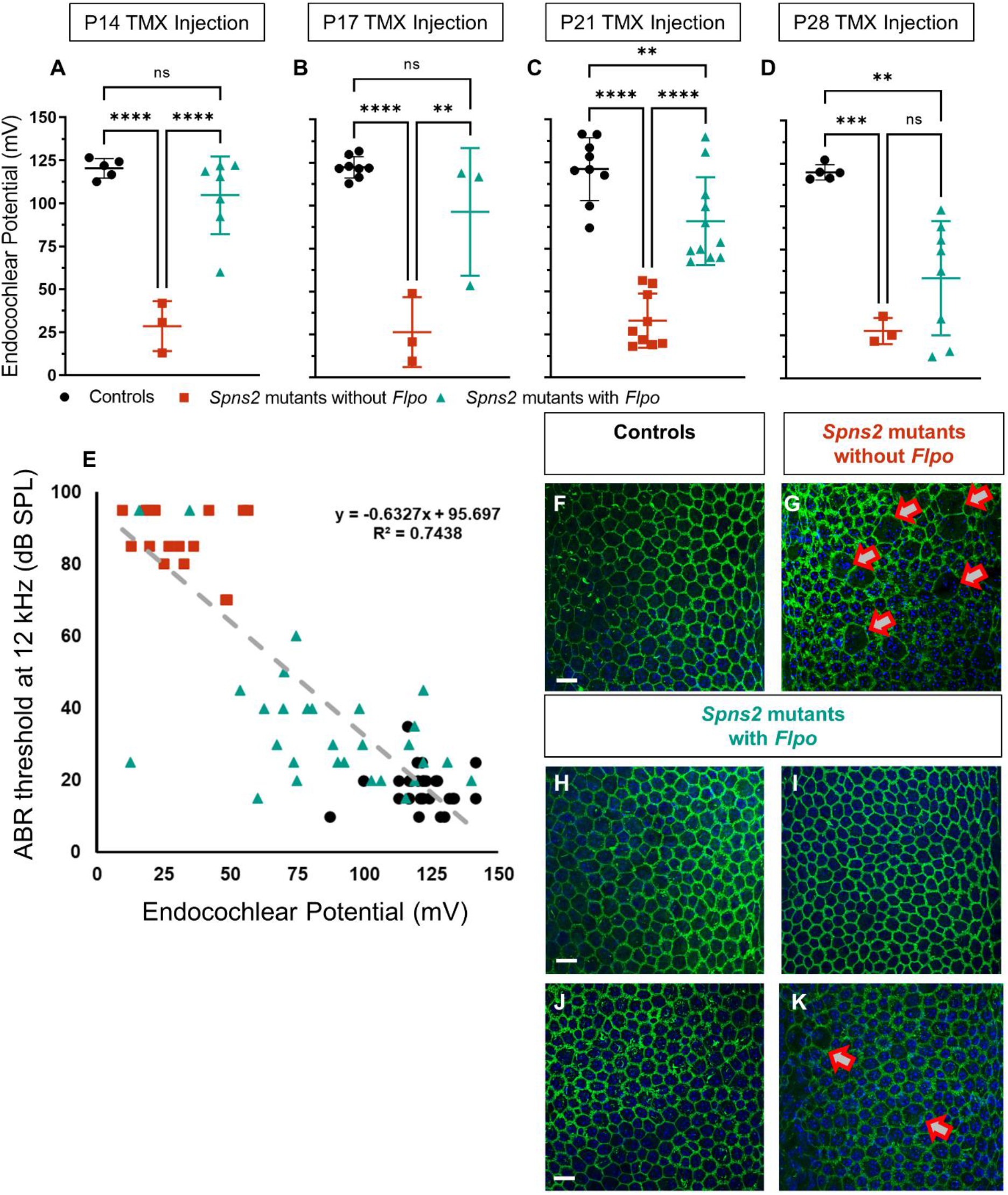
**A-D**. In the mice injected at P14 (A) there was no significant difference in EP between the controls and *Spns2* mutants with *Flpo*, while the *Spns2* mutants without *Flpo* showed a reduced EP. Controls (n = 5) 120.4 mV ± 5.7, *Spns2* mutants without *Flpo* (n = 3) 28.6 mV ± 14.6, *Spns2* mutants with *Flpo* (n = 7) 104.7 mV ± 22.7. In the mice injected at P17 (B) there was no significant difference in EP between the controls and *Spns2* mutants with *Flpo*, while the *Spns2* mutants without *Flpo* showed a reduced EP. Controls (n = 8) 122.3 mV ± 6.1, *Spns2* mutants without *Flpo* (n = 3) 26.4 mV ± 20.3, *Spns2* mutants with *Flpo* (n = 3) 96.3 mV ± 37.1. In the mice injected at P21 (C), the EP is reduced in both *Spns2* mutants with *Flpo* and *Spns2* mutants without *Flpo* in comparison with littermate controls. However, the EP in *Spns2* mutants with *Flpo* is higher than *Spns2* mutants without *Flpo*. Controls (n = 9) 121.3 mV ± 18.2, *Spns2* without *Flpo* (n = 9) 33.1 mV ± 15.8, *Spns2* mice with *Flpo* (n = 11) 90.9 mV ± 25.4. In the mice injected at P28 (D), the EP was reduced in both *Spns2* mutants with *Flpo* and *Spns2* mutants without *Flpo* in comparison with littermate controls. Controls (n = 5) 119.9 mV ± 4.4, *Spns2* mutants without *Flpo* (n = 3) 58.3 mV ± 33.2, *Spns2* mutants with *Flpo* (n = 8) 27.7 mV ± 7.6. Single mice are plotted as black circles (controls), red squares (*Spns2* mutants without *Flpo*) and teal triangles (*Spns2* mutants with *Flpo*) and as mean ± standard deviation. One-way ANOVA test with Tukey post-hoc, ** p ≤ 0.01, *** p ≤ 0.001, **** p ≤ 0.0001, ns= not significant. **E**. With an R^2^ value of 0.74, a correlation between a lower ABR threshold at 12 kHz and higher endocochlear potential was observed. All of the mice injected at P14, P17, P21, and P28 are plotted in this graph. On the y-axis are their ABR threshold values at 12 kHz at 8 weeks old, and on the x-axis is their EP measured at 8 weeks old. Controls in black circles (n = 27), *Spns2* mutants without *Flpo* in red squares (n = 18), and *Spns2* mutants with *Flpo* in teal triangles (n = 29). **F-K**. Strial marginal cell boundaries show normal morphology in *Spns2* mutants with *Flpo* injected at P14 (H),P17 (I) and P21 (J) like the controls (F, P17 injected mice), while abnormalities were present in *Spns2* mutants without *Flpo* as indicated by the red arrows (G, P17 injected mice). *Spns2* mutants with *Flpo* injected at P28 showed some morphological variabilities: in two of the four samples, no obvious defects were observed; however, in one sample, the marginal cell boundaries were defective, as observed in the *Spns2* mutants without *Flpo*; and in a second sample, as shown in panel K, only some of the boundaries were defective. The samples used for this histological analysis were the same used for the repeated ABR and EP measurements. Actin filament (Phalloidin) in green and nuclei (DAPI) in blue. Scale bar 20 μm. Controls n = 4 at P14, n = 3 at P17, n = 1 at P21 and n = 2 P28; *Spns2* mutants without *Flpo* n = 2 at P17, n = 3; *Spns2* mutants with *Flpo* n = 4 at P14, n = 5 at P17, n =3 and n =4.

The stria vascularis on the lateral wall of the cochlear duct generates EP by active pumping of cations into the endolymph, and in *Spns2*^*tm1a*^ homozygotes it shows disruption of the regular arrangement of marginal cell boundaries on the lumenal surface that is associated with reduced EP (Chen et al 2014). We examined whole mounts of the stria vascularis from the same tamoxifen-injected mice used for ABR and EP measurements. Actin filaments at the boundaries of marginal cells were labelled using phalloidin. Strial surface preparations from *Spns2* mutants with *Flpo* injected at either P14 or P17 or P21 (Fig. 4H, I and J) had normal morphology of the marginal cell boundaries (green) in contrast with *Spns2* mutants without *Flpo* which showed a disrupted pattern (Fig. 4B red arrows indicating the abnormal marginal cells boundaries). Morphological variability was observed in *Spns2* mutants with *Flpo* injected at P28, as some samples had normal marginal cell boundaries, while others showed some defect (Fig. 4K).

### Does hair cell degeneration limit the critical period for reversing hearing loss?

Sensory hair cell degeneration occurs in *Spns2*^*tm1a*^ homozygotes but this is not the cause of their hearing loss because ABR thresholds are already considerably raised before the onset of hair cell loss at around P28 (Chen et al 2014). However, hair cell loss might contribute to the efficiency of reversal of hearing loss when *Spns2* is activated at later ages in our study. Therefore we analysed the organ of Corti in whole mount preparations to assess hair cell loss two weeks after tamoxifen injection at either P14 or P28. In mice injected with tamoxifen at P14 there was no significant loss of either inner or outer hair cells in the *Spns2* mutants carrying *Flpo* compared with the control mice (Fig. 5A, C, G, I; compare black with teal bars), while *Spns2* mutants without *Flpo* (red bars) showed extensive loss of hair cells in the basal turn, representing the regions most sensitive to high frequencies (Fig. 5B, G, I). Tamoxifen injection at P28 was too late to avoid outer hair cell degeneration in the basal turn (Fig. 5D-F, H, J).

**Figure 5.**
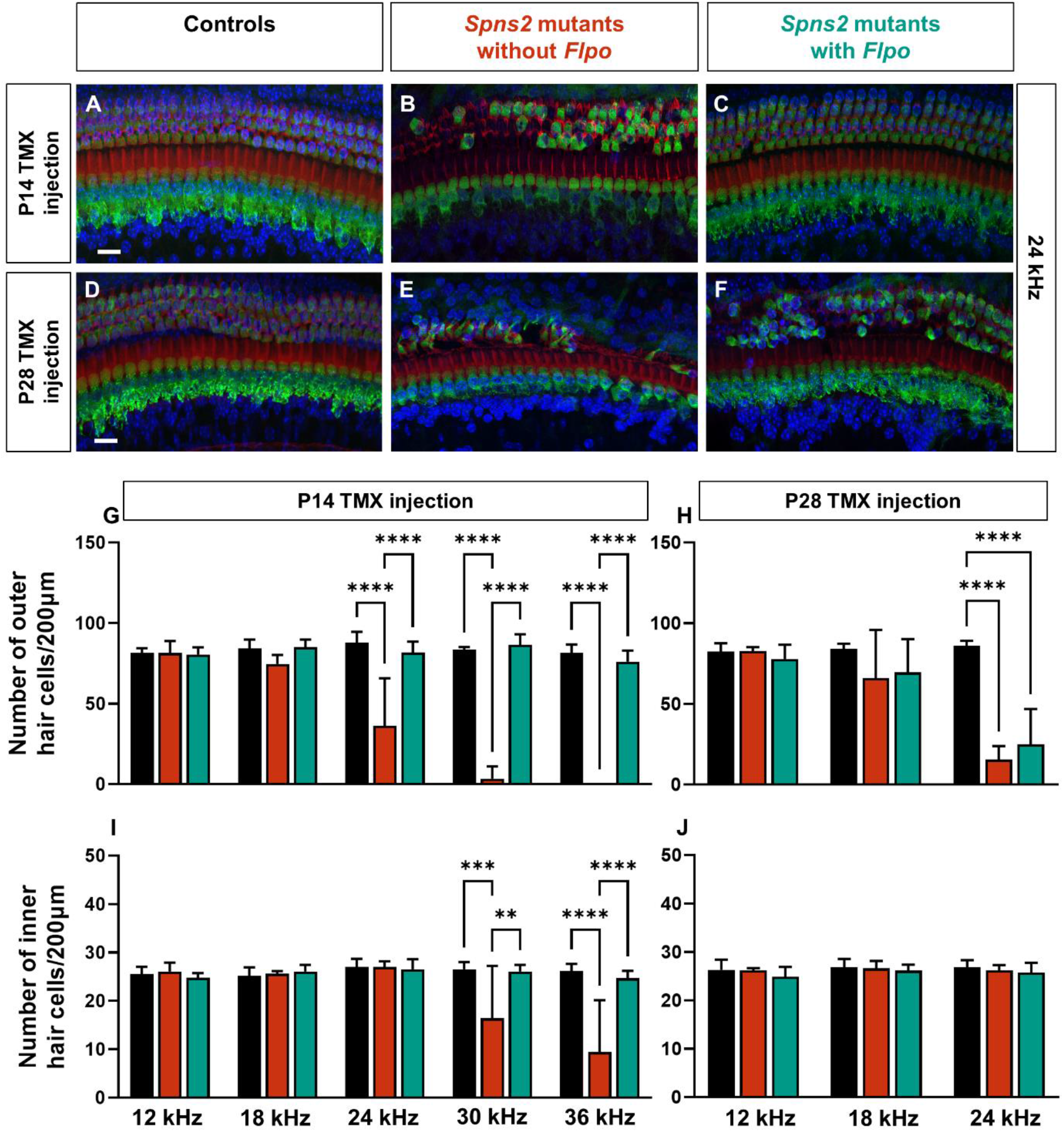
**A-F**. Surface preparation of the organ of Corti at the 24 kHz best-frequency cochlear region. Myo7a labels the hair cells in green, Phalloidin the stereocilia in red, and DAPI the nuclei in blue. Scale bar 20 μm. **A-C** and **G, I**. Tamoxifen was administered to mice at P14, and cochlear samples were collected two weeks later, at four weeks old. At all cochlear regions, the number of both outer (OHCs) and inner hair cells (IHCs) in *Spns2* mutants with *Flpo* (teal) is comparable to littermate controls (black). In contrast, *Spns2* mutants without *Flpo* (red) have significantly fewer OHCs at the 24, 30, and 36 kHz cochlear regions, as well as fewer IHCs at 30 and 36 kHz. Controls n = 6, *Spns2* mutants with *Flpo* n= 5, *Spns2* mutants without *Flpo* n = 2-5 (only at 12, 18 and 24 kHz). The cochlear duct toward the basal cochlear turn was more challenging to keep intact during the fine dissections in *Spns2* mutants without *Flpo*, so the n varies at 30 and 36 kHz. The 30 kHz cochlear region was completely missing in one of five *Spns2* mutants without *Flpo*, and the OHC area was damaged in another, so the n in panels G and I are 3 and 4, respectively. The 36 kHz cochlear region was completely absent in two of five *Spns2* mutants without *Flpo*, and only the OHC area was damaged in another, so the n in panels G and I are 2 and 3, respectively. **D-F** and **H, J**. Tamoxifen was administered to mice at P28, and cochlear samples were collected two weeks later, at six weeks old. Both *Spns2* mutants without *Flpo* and *Spns2* mutants with *Flpo* have fewer OHCs at 24 kHz in comparison with the littermate controls. Controls n = 6, *Spns2* mutants without *Flpo* n = 5, *Spns2* mutants with *Flpo* n = 7. All data are shown as mean ± SD and statistically analysed by one-way ANOVA test with Tukey post-hoc, ** p-value ≤ 0.01, *** p-value ≤ 0.001, **** p ≤ 0.0001.

## Discussion

We found that hearing impairment due to the *Spns2* mutation can be reversed after it was first detected by activating the *Spns2* gene, and ABR thresholds improve in individual mice. Neurological disorders are often thought to be irreversible, but there have been several reports that some phenotypes can be reversed in mouse models, including Rett syndrome (Guy et al 2007), Kabuki syndrome (Benjamin et al 2017), Rubinstein-Taybi syndrome (Alarcón et al 2004), *SYNGAP1* deficiency (Creson et al 2019), and autism spectrum disorders due to *SHANK3* mutations (Mei et al 2016). More recently, it was reported that mice with an *Otof* mutation can show improvements in ABR thresholds following gene therapy as late at 30 days after birth (Akil et al 2019). Our finding of reversal of hearing loss in individual mice after activation of the *Spns2* gene is an important proof-of-concept that deafness associated with EP deficiency not only can be halted in its progression but also can be reversed. This demonstration raises the prospect that this type of hearing loss may be reversible in humans.

Histopathology suggests that human age-related hearing loss can affect three main sites within the cochlea: sensory hair cells; synapses between hair cells and cochlear neurons; and the stria vascularis on the lateral wall of the cochlear duct (Schuknecht & Gacek 1993). Analysis of human audiogram shapes compared with experimental damage to either the stria vascularis or hair cells supports the concept that strial dysfunction has a role to play in human hearing loss (Dubno et al 2013; Vaden et al 2022). Furthermore, recent meta-analysis of genome-wide association studies of hearing loss in the human population provides additional support for the importance of strial function in the pathogenesis of hearing loss (Trpchevska et al 2022). Thus, this type of pathology is a good target for development of generic treatments to boost strial function.

We discovered that the earlier the activation of the gene, the better the restoration of cochlear function, including ABR thresholds and waveforms, EP levels and hair cell preservation. This suggests that there is a critical period for reversing hearing loss, with the precise timing of this period depending upon the outcome measure. For example, *Spns2* activation at P17 was effective at reversing raised ABR thresholds at frequencies of 18kHz and below but was too late to rescue thresholds at higher frequencies, 24kHz and above. The loss of hair cells when *Spns2* is activated later is likely to influence the endpoint of the critical period for reversing hearing loss. However, the earliest electrophysiological deficit recorded was the reduced EP (Chen et al 2014), and we found that later activation of *Spns2* led to lower EP levels (Fig. 4A-D). Although we cannot directly record a reversal in EP level in an individual mouse in the days following tamoxifen injection, our observations suggest that the EP can probably be increased by activating *Spns2* and that it is likely that ABR thresholds reflect this. The finding of a critical period indicates that early intervention in humans affected by the strial type of pathology would be important, before the changes leading to EP reduction become irreversible and before secondary hair cell degeneration occurs.

In summary, our finding is a proof-of-concept that at least one type of hearing loss can be reversed. Furthermore, the availability of over 4,000 mouse mutants carrying this design of mutation, many of which show abnormal phenotypes (https://www.i-dcc.org/imits; https://www.mousephenotype.org/; Birling et al 2021; Groza et al 2023), facilitates similar investigations in other diseases that up to now have been considered irreversible.

## Materials and Methods

### Ethics statement

Mouse studies were carried out in accordance with UK Home Office regulations and the UK Animals (Scientific Procedures) Act of 1986 (ASPA) under UK Home Office licences, and the study was approved by the King’s College London Ethical Review Committees. Mice were culled using methods approved under these licences to minimize any possibility of suffering.

### Generation of mutant mice

*Spns2*^*tm1a(KOMP)Wtsi*^ (*Spns2*^*tm1a*^) mutant mice were generated at the Wellcome Trust Sanger Institute on a C57BL/6N genetic background (White et al. 2013, Chen et al. 2014). B6N.129S6(Cg)-*Gt(ROSA)26Sor*^*tm3(CAG-flpo/ERT2)Alj*^/J mice, also known as *R26*^*FlpoER*^ mice, express a tamoxifen-inducible, optimized FLPe recombinase variant called Flpo and were obtained from the Jackson Laboratory (Lao et al 2012). Both mutant lines are available from public repositories (EMMA and The Jackson Laboratory). We crossed mice carrying the *Spns2*^*tm1a*^ and *R26*^*FlpoER*^ alleles to generate littermates including the genotypes for analysis: *Spns2*^*+/+*^ and *Spns2*^*+/tm1a*^ with and without *Flpo* which formed the normal control group; *Spns2*^*tm1a/tm1a*^ homozygotes without *Flpo*, providing the mutant group; and *Spns2*^*tm1a/tm1a*^ homozygotes with *Flpo*, which were the experimental mice. All genotypes were injected with tamoxifen (see below).

### Genotyping

DNA was extracted from pinna skin and used as the template for short range PCR using a common *Spns2* forward primer (5′ CAAAACAATATGGGCTGGGG3′), a *Spns2* reverse primer (5′ GATGAAGGCAGGACTCAGGG3′) and a mutant-specific reverse primer (5′TCGTGGTATCGTTATGCGCC3′) (Fig.1 A, blue and black arrows). The resulting band sizes were 363 bp for the *Spns2* wildtype product and 187 bp for the mutant product, as previously described (Chen et al., 2014). The wildtype product was not amplified from the *tm1a* mutant allele because the primers bound to sites that were too far apart for short-range PCR to be successful, but it can be amplified in the *tm1c* allele (Fig. 1). Upon tamoxifen injection, Flpo recombinase is tranlocated to the nucleus where it recognises the FRT sites in the *Spns2*^*tm1a*^ allele, inducing recombination with the consequent removal of the mutagenic cassette and restoration of *Spns2* gene activity. The allele generated from this recombination is called *Spns2*^*tm1c*^. The native WT *Spns2* allele had a 363bp PCR product, while the *Spns2*^*tm1c*^ allele had a 463bp PCR product using the same primers (Fig1 A), 100bp larger due to the presence of one FRT site and one LoxP site.. The presence of the *Flpo* allele was checked using the common Rosa26 forward primer 5’AAAGTCGCTCTGAGTTGTTAT3’ with either the Rosa26 reverse primer 5’GGAGCGGGGAAATGGATATG3’, or the Flpo mutant reverse primer 5’ TTATGTAACGCGGAACTCCA3’ (Fig.1 A, magenta and grey arrows) The resulting band sizes were 603 bp for the wildtype product and 309 bp for the mutant product.

### Tamoxifen-induced gene recombination

Tamoxifen in corn oil (Sigma-Aldrich, C8267) was injected intraperitoneally using a single injection at a dose of 0.2mg/g at one of four different ages: postnatal day (P)14, P17, P21 or P28 (Fig. 1B). Auditory Brainstem Responses (ABRs) were recorded the same day and before the tamoxifen injection and then repeated ABR tests were performed up to 8 weeks old (Fig.2) when the EP was also measured as a terminal procedure (Fig. 4 A-D). The Flpo-FRT recombination was verified on pinna tissue upon DNA extraction and PCR testing using the same *Spns2* primers used for genotyping of the *Spns2* allele. Our results showed the expected recombination and generation of the *Spns2*^*tm1c*^ allele (Fig 1B).

### Auditory Brainstem Response (ABR) recording

Mice were anaesthetised by intra-peritoneal injection of 100 mg/kg Ketamine (Ketaset, Fort Dodge Animal Health) and 10 mg/kg Xylazine (Rompun, Bayer Animal Health). Brainstem auditory evoked potentials were measured in a sound-attenuating chamber fitted with a heating blanket as previously described (Ingham et al. 2011; Ingham 2019). Subcutaneous recording needle electrodes (NeuroDart; Unimed Electrode Supplies Ltd, UK) were inserted on the vertex and overlying the left and right bullae. Responses were recorded to free-field calibrated broadband click stimuli (10 μs duration) and tone pips (5 ms duration, 1 ms onset/offset ramp, fixed phase onset) at frequencies between 3 and 42 kHz, at levels ranging from 0-95 dB SPL in 5 dB steps, at a rate of 42.6 stimuli per second. Evoked responses were digitized, amplified and bandpass filtered between 300-3000 Hz, and 256 responses averaged for each frequency/intensity combination to give a waveform. ABR thresholds were defined as the lowest stimulus level to evoke a visually-detected waveform.

### Endocochlear Potential (EP) recordings

Mice were anaesthetised with intra-peritoneal urethane (0.1 mL/10g body weight of a 20% w/v solution of urethane) and EP was measured as described previously (Steel and Barkway, 1989; Ingham et al., 2016). A reference electrode (Ag-AgCl pellet) was placed under the skin of the neck. A small hole was made in the basal turn lateral wall and the tip of a 150 mM KCl-filled glass micropipette was inserted into the scala media. The EP was recorded as a stable positive potential compared with the reference electrode.

### Hair cell quantification, lateral wall surface preparation and confocal imaging

The organ of Corti was analysed in mice injected with tamoxifen at P14 and P28 and then collected 2 weeks after tamoxifen injection (Fig.5). The lateral wall surface preparation was performed using the inner ears of the mice that were injected with tamoxifen and had the repeated ABR tests (Fig. 2) as well as the EP measurements recorded (Fig. 3). Inner ears were fixed in 4% paraformaldehyde for 2 hours and decalcified in EDTA overnight at room temperature. Following fine dissection, both the lateral wall and the organ of Corti specimen were permeabilised in 5% Tween PBS for 40 minutes and incubated in blocking solution (4.5 mL of 0.5% Triton X-100 in PBS and 0.5 mL of normal horse serum) for 2 hours. The organ of Corti samples were incubated in Myo7a primary antibody (diluted 1:200; 25-6790, Proteus) overnight at 4°C and thenincubated for 45 minutes at room temperature with the secondary antibody goat anti-rabbit IgG Alexa Fluor488 (1:300, #A11008, Thermo Fisher Scientific) and phallodin (1:500, #R415, Thermo Fisher Scientific). The lateral wall samples were incubated with phalloidin for 45 minutes at room-temperature. Both the lateral wall and organ of Corti specimen were mounted using ProLong Gold mounting media with DAPI (P36931, Life Technologies) and stored at 4°C. Specimens were imaged using a Zeiss

Imager 710 confocal microscope interfaced with ZEN 2010 software. The plan-APOCHROMAT 40x Oil DIC objective was used and brightness and contrast were normalised for the dynamic range in all images. Z-stacks were collected at 0.5μm and maximum intensity projection images were generated for analysis. The best-frequency areas were determined according to the mouse tonotopic cochlear map using the ImageJ plugin “Measure Line” (Hickman et al., 2018; Mueller et al 2005). Manual quantification of both outer and inner hair cells (OHCs and IHCs) was performed at specific frequency regions.

## Acknowledgements

We thank the Wellcome Sanger Institute Mouse Genetics Project for generating and providing the *Spns2*^*tm1a*^ mutant mouse.

## Data availability

All data are provided within the report and the two mutant mouse lines used are available from public repositories.

## Author contributions

EM, NJI and KPS designed the research; EM and NJI carried out the experiments; KPS obtained the funding; EM, NJI and KPS analysed the data; EM, NJI and KPS wrote the paper.

The authors declare no competing interest.

This report will be published as an open access contribution with a CC/BY licence.

